# Pandemic Publishing: Medical journals drastically speed up their publication process for Covid-19

**DOI:** 10.1101/2020.04.18.045963

**Authors:** Serge P.J.M. Horbach

## Abstract

In times of public crises, including the current Covid-19 pandemic, rapid dissemination of relevant scientific knowledge is of paramount importance. The duration of scholarly journals’ publication process is one of the main factors hindering quick delivery of new information. While proper editorial assessment and peer review obviously require some time, turnaround times for medical journals can be up to several months, which is undesirable in the era of a crisis. Following initiatives of medical journals and scholarly publishers to accelerate their publication process, this study assesses whether medical journals have indeed managed to speed up their publication process for Covid-19 related articles. It studies the duration of 14 medical journals’ publication process both during and prior to the current pandemic. Assessing a total of 669 articles, the study concludes that medical journals have indeed drastically accelerated the publication process for Covid-19 related articles since the outbreak of the pandemic. Compared to articles published in the same journals before the pandemic, turnaround times have decreased on average by 49%. The largest decrease in number of days between submission and publication of articles was due to a decrease in the number of days required for peer review. For articles not related to Covid-19, no acceleration of the publication process is found. While the acceleration of journals’ publication process is laudable from the perspective of quick information dissemination, it also raises concerns relating to the quality of the peer review process and the quality of the resulting publications.

## Background

The world is facing an unprecedented health crisis affecting nearly all parts of society. In these times, access to the most state-of-the-art scientific knowledge is paramount to tackling the crisis. Academic journals and scholarly publishers are hence called upon to make new knowledge openly available and deliver new insights quickly.

In the current Covid-19 era, it is clear that new knowledge is direly needed. Scientists all over the world have stepped in to do experiments, observational studies and new analyses as to obtain relevant information. However fast we would like to have access to this information, the scientific method used to obtain it, requires time. Drug trials and vaccine creations do not happen overnight (Thorp, 2020). However, once such information has been gathered, it needs to be disseminated to all those potentially in the position to use it, as quick as possible. Traditionally, scholarly journals have been one of the main outlets to facilitate this (Horbach & Halffman, 2018).

One of the factors hindering quick delivery of new information through scholarly journals is the duration of their publication process. Through editorial assessment and peer review, journals select which articles deserve to be published on their pages, ideally filtering out invalid, erroneous or otherwise problematic research. Even though celebrated as being one of the hallmarks of science, the editorial process is also regularly criticized. Commentators blame it for being inconsistent (Peters & Ceci, 1982), essentially flawed (Smith, 2006), biased (Teplitskiy, Acuna, Elamrani-Raoult, Körding, & Evans, 2018), and – particularly relevant in these times of crisis – slow (Nguyen et al., 2015).

Several studies have previously aimed to assess the typical duration of journals’ publication process (Lin, Hou, & Wu, 2016; Tosi, 2009). In their analyses, researchers commonly distinguish two stages of this process: the review stage (i.e. the stage between article submission and formal acceptance) and the editorial stage (i.e. the stage between acceptance and final publication, either online or in print). In a meta-analysis including over 2700 journal articles, Björk and Solomon (2013) find considerable differences in turnaround times (i.e. the period between submission and publication of a journal article, sometimes also called ‘publication delays’) between research disciplines. For biomedical journals, they find an average duration of the review stage of just over four months, while the editorial stage takes on average about five months. Clearly, such turnaround times are highly undesirable in light of the current health crisis.

Two major responses to circumvent long turnaround times can currently be witnessed. From an author perspective, commentators are reporting a sharp increase in the use of preprint servers. On these online platforms, authors upload their manuscript, making it publicly accessible immediately upon finalization of the text (Gunnarsdottir, 2005). Because no review, editorial assessment or copyediting takes place, manuscripts can be made accessible without publication delay. However, as manuscripts are only reviewed once they are available for anybody to read and use, scholars warn for potentially incorrect results spreading without editorial assessment filtering them. In fact, several cases of invalid research regarding Covid-19 being published as preprints have already been reported (Heimstädt, 2020; Marcus & Oransky, 2020). It should be noted though that this is not specific to preprints, as journal articles can require from post-publication corrections and retractions as well (Horbach & Halffman, 2019). Several articles related to Covid-19 have already gone through this process (Gautret et al., 2020).

From the publishers’ side, several journals and publishers are currently modifying their editorial procedures and policies to warrant fast dissemination of relevant information. For instance, *eLife* announced it would curtail requests for additional experiments during revisions, suspend its deadline for submitting revisions, make the posting of preprints to bioRxiv or medRxiv the default for all eLife submissions, and it would specifically mobilize early-career researchers to become reviewing editors and reviewers in order to extent the journal’s reviewer pool (Eisen, Akhmanova, Behrens, & Weigel, 2020). Similarly, Nature put out an open invitation to researchers with relevant expertise to review Covid-19 related papers over short time (“Coronavirus pandemic: Nature’s pledge to you,” 2020). Hence, journals and publishers are aiming to attract reviewers that can assist in the rapid publication of new findings, relevant to tackle the health crisis. The *Medical Journal of Australia (MJA)* has drafted policies related to both preprints are rapid peer review, setting up ‘fast lanes’ for Covid-19 related research:

> “The *MJA* has stepped up to play its part in meeting this crisis, including ultra-rapid review of SARS-CoV-2 manuscripts and pre-print publication of unedited papers, to ensure that the newest data and viewpoints are available as soon as possible.”
>
> — (Talley, 2020)

The *Royal Society Open Publishing* announced to establish a similar fast lane for their registered reports on Covid-19 related content. They have even gathered a group of 700 reviewers who have committed to review a paper in 24 to 48 hours when called up on (Brock, 2020). The journal also acknowledges one of the major concerns related to these fast dissemination models: “The ultra-rapid review and publication model entails a risk of error, but sharing important information too slowly is a much greater hazard.” (Talley, 2020)

In this article, we assess whether the scholarly publishing community succeeds in speeding up the dissemination of Covid-19 related content. To do so, we assess the use of preprint servers, the uptake of preprint articles in academic journals, and the duration of journals’ publication process both prior to and during the present pandemic.

## Methods

For our analysis, we use a repository of Covid-19 related research articles established by the Centre for Science and Technology Studies (CWTS). The repository is based on databases of CORD19, Dimensions and the World Health Organisation (WHO), and includes articles on Covid-19, SARS-CoV-2, and related (corona) viruses and infectious diseases. In particular, this means that the database contains journal articles and preprints that predate the current Covid-19 pandemic, as it for instance also includes articles on the 2002 SARS virus and disease. For brevity’s sake, all such articles will in the remainder of this article by described as ‘Covid-19 related’ articles. A full description of the database as well as access to all relevant data is available (CWTS, 2020). Colavizza et al. (2020) provide a description and analysis of parts of this database. All results in this article are based on the April 4 release of the database. We note that the majority of articles in the dataset originates from the CORD19 database. Some doubt has been raised about the relevance of some of this database’s articles to the current pandemic (Colavizza et al., 2020), but for our purposes, the scope of the database seems reasonable. Based on this dataset, several analyses were performed:

### Duration of journals’ publication process

We analysed the duration of the publication process, in number of days, for a sample of 529 journal articles. 259 articles were published during the present pandemic (i.e. from Jan 1, 2020) and 270 were published prior to the pandemic (i.e. before Oct 1, 2019). The articles were published in 14 different journals. Journals were selected based on their number of articles both prior to and during the current pandemic as well as the availability of data on when articles were submitted, accepted and published. We selected the ten journals publishing most Covid-19 related articles in general, supplemented by the five journals publishing most Covid-19 related content since the start of the pandemic, that make publication data (submission, acceptance and publish) available. One journal, *Viruses,* matches both criteria. The list of journals used in this analysis, including their number of articles, as well as the journals discarded because no data on submission, acceptance or publication dates was available, is added as supplementary material A. From those journals, we sampled all articles published since the start of the pandemic and matched those to an equal number of articles published in the same journal prior to the pandemic to form a control group. In case the control group had fewer articles, we used this number of articles and only selected the most recent articles after the pandemic. In case less than ten articles were published since the start of the pandemic, we nonetheless sampled ten articles for the control group. For the control group, we sampled articles starting with publications in 2019 (but before Oct 1^st^) and moving backwards, in order to make sure editorial policies most closely resemble those in the pandemic. Table 1 presents the list of journals used, including the number of articles sampled per journal. Information on the dates of submission, acceptance and publication was manually retrieved from the journal’s webpage. In case journals distinguished between publication online and appearance in the print issue of the journal, we selected the date of online publication.

**Table 1.** duration of the publication process for Covid-19 related papers, distributed over Review stage (between submission and acceptance) and Editorial stage (between acceptance and publication). Data distinguishes between the period before and during the Covid-19 pandemic. All durations are given in days.

|  | Number of articles |  |  | Number of days for publication - During Pandemic |  |  |  |  | Number of days for publication - Before Pandemic |  |  |  |  |
| --- | --- | --- | --- | --- | --- | --- | --- | --- | --- | --- | --- | --- | --- |
| Journal title | During Pandemic | Before Pandemic | Total | Review Stage | Editorial Stage | Entire Publication Process | 95% CI Entire publication process |  | Review Stage | Editorial Stage | Entire Publication Process | 95% CI Entire publication process |  |
| Archives of Virology | 9 | 10 | 19 | 127.1 | 70.1 | 197.2 | 162.0 | 232.4 | 122.6 | 37.9 | 160.5 | 107.0 | 214.0 |
| Eurosurveillance | 30 | 30 | 60 | 8.3 | 1.7 | 10.0 | 7.1 | 12.9 | 105.9 | 62.4 | 168.3 | 113.2 | 223.5 |
| International Journal of Infectious Diseases | 32 | 32 | 64 | 23.7 | 5.3 | 28.9 | 14.8 | 43.1 | 77.0 | 8.5 | 85.5 | 64.7 | 106.3 |
| Journal of Hospital Infection | 24 | 24 | 48 | 10.0 | 5.6 | 15.6 | 1.8 | 29.5 | 59.5 | 15.8 | 75.3 | 55.9 | 94.7 |
| Journal of Medical Virology | 13 | 13 | 26 | 10.3 | 4.9 | 15.2 | 11.8 | 18.7 | 107.3 | 64.4 | 117.3 | 56.0 | 178.7 |
| Journal of Virology | 4 | 10 | 14 | 37.0 | 41.5 | 78.5 | 19.6 | 137.4 | 61.5 | 77.9 | 139.4 | 119.5 | 159.3 |
| PLoS ONE | 14 | 14 | 28 | 183.3 | 25.9 | 209.2 | 153.9 | 264.5 | 230.9 | 25.9 | 256.7 | 164.2 | 349.2 |
| Scientific Reports | 14 | 14 | 28 | 147.9 | 18.4 | 166.4 | 133.0 | 199.7 | 216.7 | 57.9 | 274.6 | 197.2 | 352.1 |
| Travel Medicine and Infectious Disease | 45 | 45 | 90 | 12.6 | 2.7 | 15.3 | 8.8 | 21.9 | 91.5 | 5.1 | 96.6 | 71.9 | 121.4 |
| Vaccine | 9 | 10 | 19 | 148.2 | 22.7 | 170.9 | 114.0 | 227.7 | 154.3 | 14.5 | 168.8 | 96.3 | 241.3 |
| Veterinary Microbiology | 9 | 10 | 19 | 89.6 | 4.3 | 93.9 | 70.5 | 117.3 | 69.2 | 2.4 | 71.6 | 59.2 | 84.0 |
| Virology | 8 | 10 | 18 | 90.0 | 3.3 | 93.3 | 63.9 | 122.6 | 63.3 | 5.6 | 68.9 | 51.3 | 86.5 |
| Virus Research | 10 | 10 | 20 | 124.2 | 2.4 | 126.6 | 89.7 | 163.5 | 119.4 | 9.2 | 128.6 | 37.4 | 219.8 |
| Viruses | 38 | 38 | 76 | 32.5 | 4.2 | 36.7 | 28.6 | 44.8 | 33.3 | 3.7 | 37.0 | 30.1 | 43.9 |
| Total | 259 | 270 | 529 | 51.0 | 9.3 | 60.3 | 50.9 | 69.8 | 95.9 | 23.6 | 117.4 | 103.9 | 130.9 |

To control for potential effects specific to Covid-19 related papers, we selected, for all journals in our sample, the ten most recently published articles (as of April 16^th^, 2020) about non-Covid-19 related content. In particular, these were articles not present in our previous dataset and articles not mentioning Covid-19, coronavirus, SARS-CoV-2, or Cov-19 in their title, keywords and abstract. All the 140 articles in this control group were published during the current pandemic, with 64% published in April 2020, 32% in March, 1% in February, and 3% in January.

### Publication of preprints

We assessed the usage of preprint servers as a fast way of disseminating academic knowledge by counting the number of preprints on Covid-19 related content both during and before the current pandemic. For this we used all preprints in de database, and did use a more narrow sampling strategy. Hence, we include all Covid-19 related preprints. In addition, we analysed the number of preprints that also appeared as journal articles, and the average number of days between publication of the preprint and the corresponding journal article. The analysis is based on the linkage of preprints and journal articles in the Dimensions Database (https://docs.dimensions.ai/dsl/index.html).

Figure 1 presents an overview of the number of journal articles and preprints per year in our dataset. Unsurprisingly, it shows a sharp increase for both publication types since the outbreak of the current pandemic. However, earlier pandemics, such as the SARS outbreak in 2002, are clearly visible as well.

**Figure 1.**
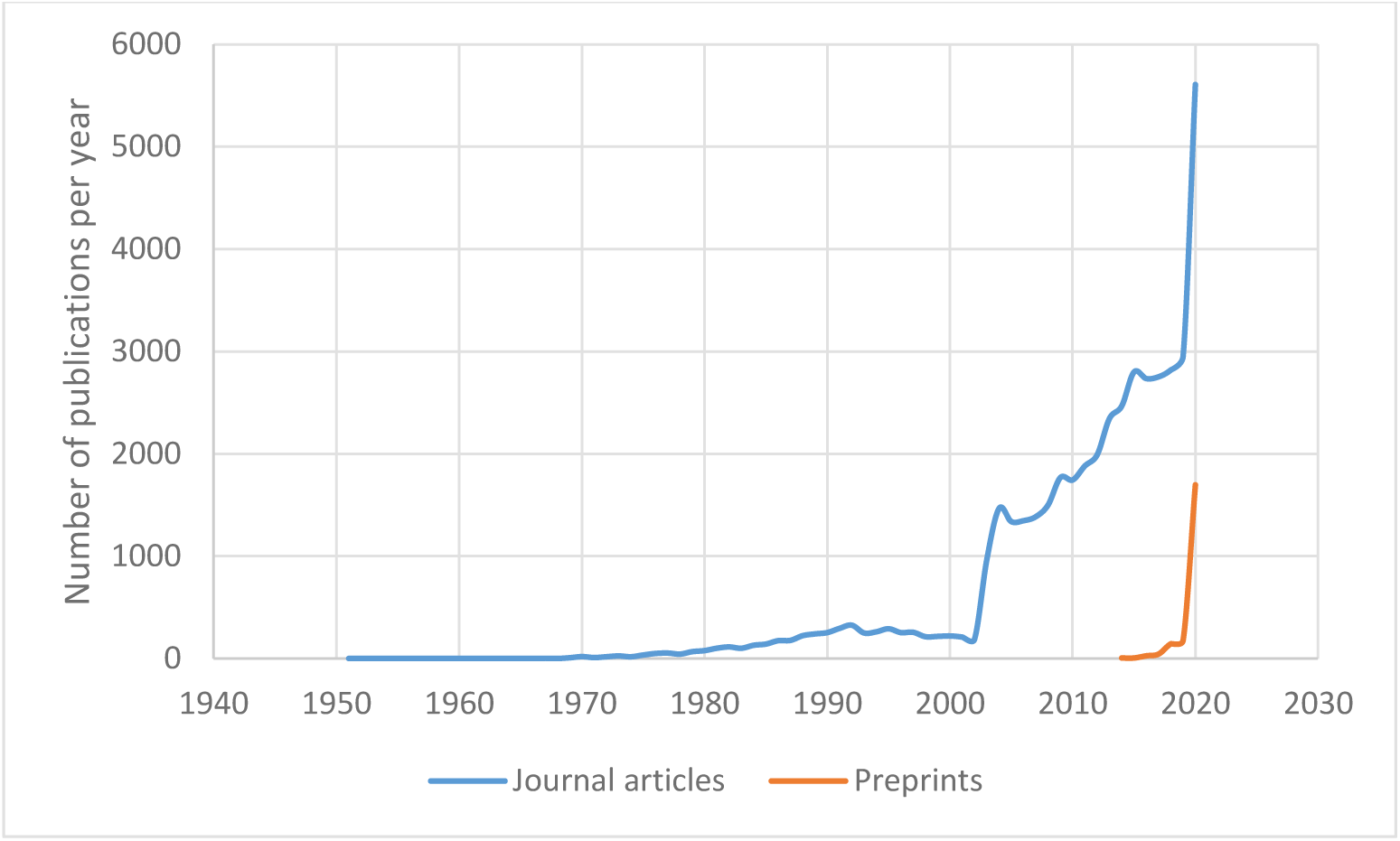
the number of journal articles and preprints related to Covid-19 per year in our database.

## Results

Figure 2 compares the overall duration of journals’ publication process prior to and during the present pandemic. It demonstrates that, on average, journals have drastically increased the speed of their processes for Covid-19 publications: average turnaround times in our journal sample has decreased from 117 to 60 days. Comparing the 95% confidence intervals of both statistics, shows the decrease to be highly substantial and significant.

**Figure 2.**
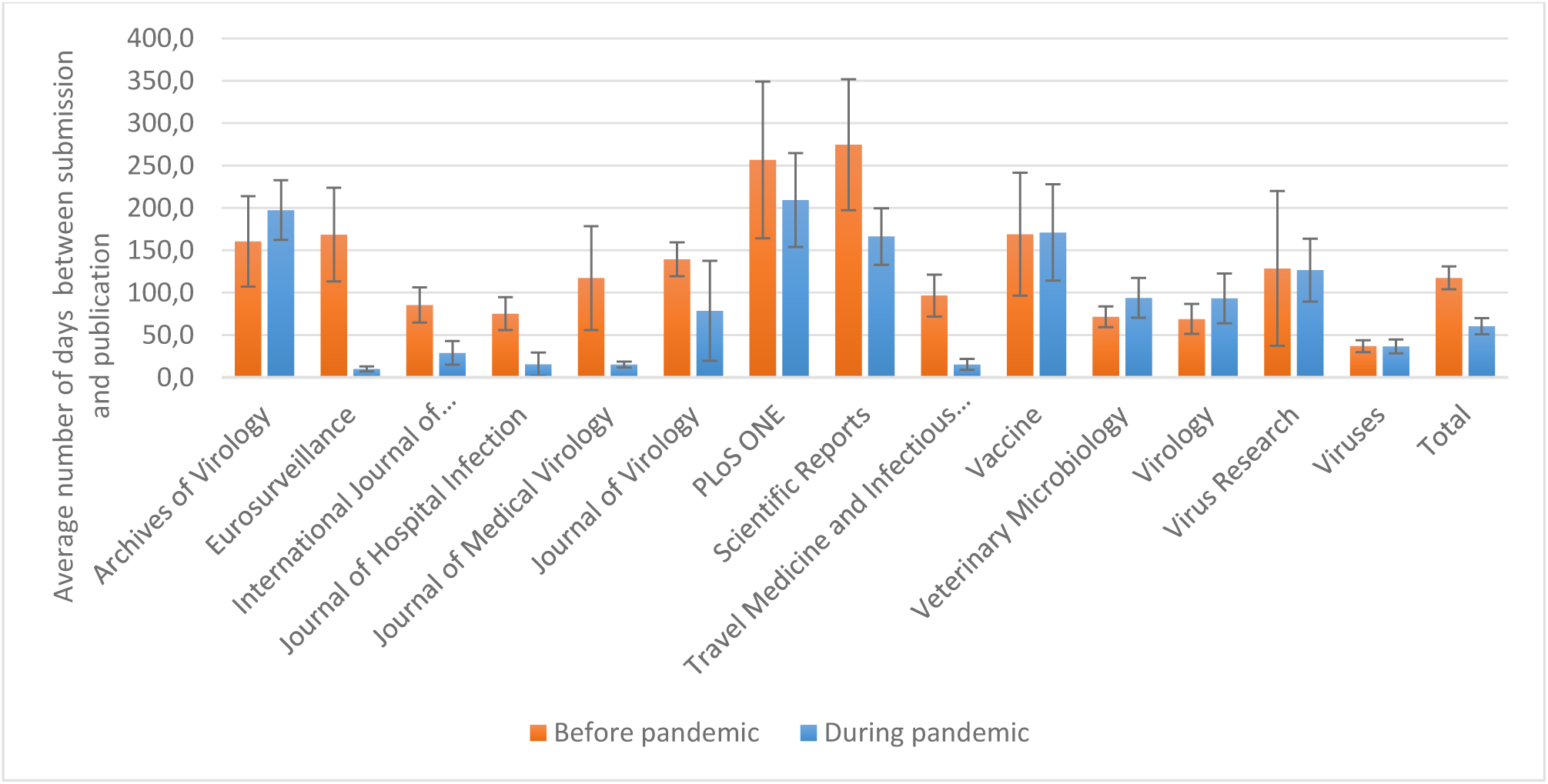
Average duration of the publication process for Covid-19 related papers in our sample of journals, as well as the total average over all journals (Total). Durations are given in number of days and distinguish between the period before and during the Covid-19 pandemic. Error bars represent the 95% confidence interval.

Table 1 presents descriptive statistics on the average duration of the publication process for Covid-19 related articles in journals in our sample. It distinguishes between the periods before and during the pandemic and it splits the entire publication process in the *Review* stage (between submission and acceptance) and *Editorial* stage (between acceptance and submission).

Figure 3 presents a graphical overview of the average decrease in turnaround time in the period during the crisis compared to the period prior to the pandemic. It again distinguishes between the Review and Editorial stages of the publication process. Note that negative numbers in this case indicate an *increase* in turnaround time. The figure indicates the Review stage shortens for ten out of the fourteen journals in our sample, while nine journals managed to shorten their Editorial stage. Average acceleration is around 50% for both stages, but it goes up to nearly 100% in some journals.

**Figure 3.**
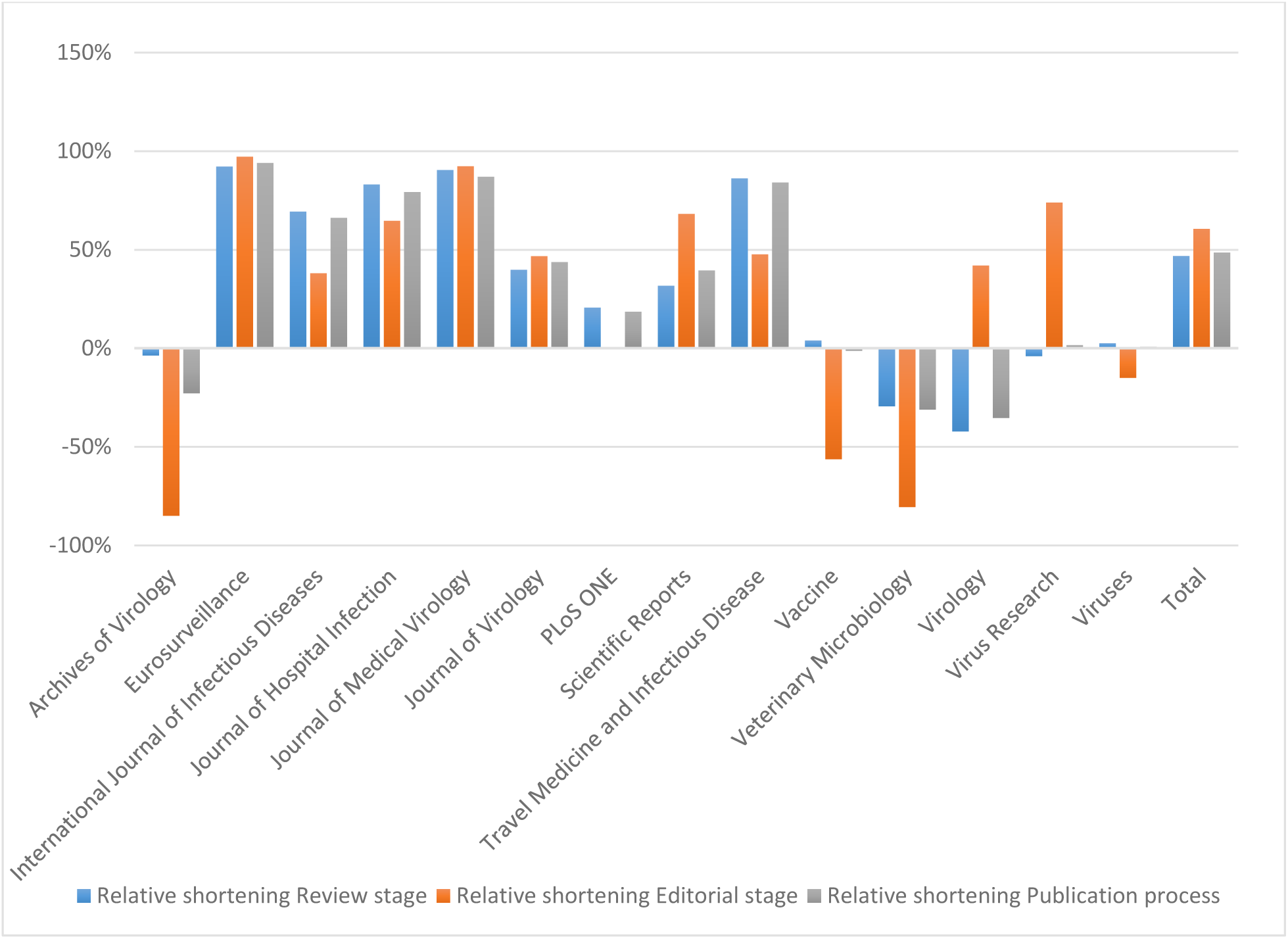
Relative average decrease in time spend per journal on the Review stage, Editorial stage and Entire publication process in the journals of our sample, as well as for the total set of journals (Total). Decrease is measured as the shortening of duration in the pandemic era compared to the pre-pandemic era, i.e. negative values indicate an increase in duration during the pandemic compared to the period before the pandemic.

To check whether the acceleration of publication processes is specific to Covid-19 related papers, we analysed the turnaround times for non-Covid-19 related articles published since the start of the pandemic. For all journals in our sample, we selected the ten most recently published articles (as of April 16^th^, 2020) about non-Covid-19 related content. In particular, these were articles not present in our previous dataset and articles not mentioning Covid-19, coronavirus, SARS-CoV-2, or Cov-19 in their title, keywords and abstract. For these articles we also analysed the turnaround times of their publication process. Results are presented in figure 4. The figure indicates that for most journals, articles not related to Covid-19 have very similar turnaround times as articles published before the pandemic. Unpacking the publication process in the Review and Editorial stage, we conclude that, again, non-Covid-19 related articles follow a very similar pattern to articles published before the pandemic.

**Figure 4.**
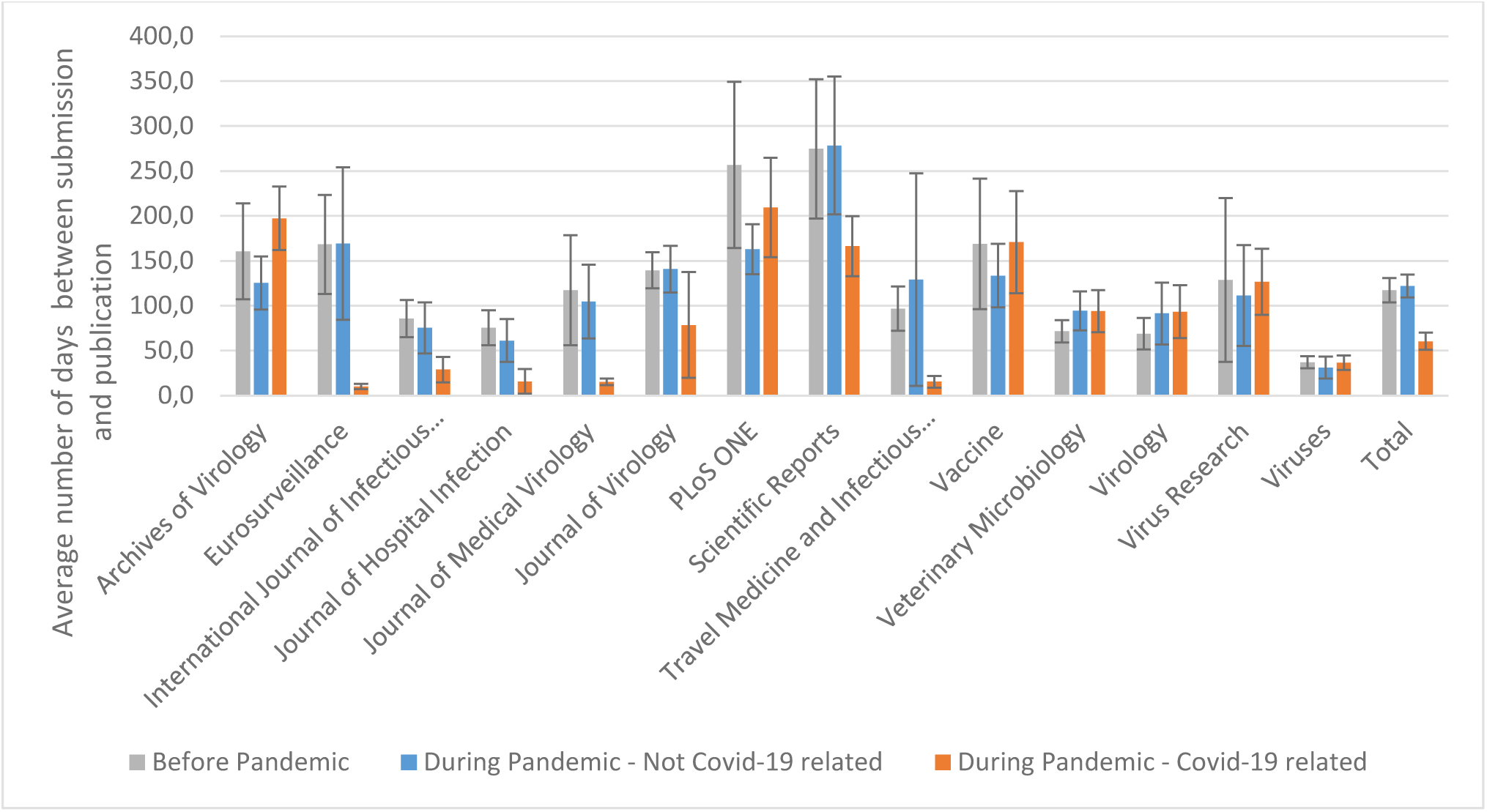
Average duration of the publication process for papers in our sample of journals, as well as the total average over all journals (Total). Durations are given for articles published before the pandemic, and articles published during the pandemic, both those related to Covid-19 and those not related to Covid-19. Durations are given in number of days. Error bars represent the 95% confidence interval.

Some of the most prominent, high impact medical journals are not part of our sample, because they do not share all relevant data on submission, acceptance and publication dates. Comparing the total number of published articles in high impact journals such as BMJ, The Lancet, JAMA and NEJM does not give a clear indication of faster publication: These four journals published 864, 421, 351, and 307 articles respectively in 2020, according to a Web of Science search. Over the same period in 2019 they published 874, 497, 335, and 334 articles respectively. Hence, most show a small decrease in the total number of published articles. Consequently, if they managed to speed up their publication process for Covid-19 related articles, this has gone at the expense of other content being published less, or less quickly.

Next we turn our analysis towards the publication of preprints. As was shown in figure 1, the number of preprints on Covid-19 related content has seen a sharp increase since the outbreak of the pandemic. In total 2102 preprints were published on seven preprint servers: SSRN Electronic Journal, bioRxiv, ChemRxiv, JMIR Preprints, Research Square, and medRxiv. We note that even though arXiv publications are included in the Dimensions database, they are not included in the 4 April release of the dataset we used, due to technical issues. Out of the 2102 preprints in our dataset, 129 have currently also appeared as journal article. Due to the small number of preprints appearing as journal articles, no statistically relevant conclusions can be drawn about the uptake of preprints in journals. However, analyse the average duration between the publication of the preprint and the corresponding journal article we see a steady increase, ranging from, on average, 137 days in 2017 and just over 200 days in 2020. Currently, we do not see any indication of acceleration of the uptake of preprints in journals since the outbreak of the current pandemic. However, the sample is small and these statements should hence be treated with caution.

The 129 preprints were published in 68 different academic journals with only five journals publishing at least five preprints. The three journals publishing most preprints were *Journal of Virology, PLoS ONE,* and *Scientific Reports*, all of which are in our sample of journals used for the previous analyses. Hence, for the preprints in these journals, we can analyse the turnaround times of their publication process. The results are presented in table 2.

**Table 2.** Average duration of the publication process for articles that previously appeared as preprints. Durations are given in number of days. The duration of the publication process is distributed over the Review stage (between submission and acceptance) and Editorial stage (between acceptance and publication). On top of the publication process, the table also indicates the average number of days between publication of the preprint and submission of the journal article.

| Journal title | Review Stage | Editorial Stage | Entire Publication Process | Time from preprint publication to journal submission |
| --- | --- | --- | --- | --- |
| Journal of Virology | 55.6 | 70.7 | 126.4 | 25.4 |
| PLoS ONE | 126.6 | 21.9 | 148.5 | 8.6 |
| Scientific Reports | 105.3 | 16.3 | 121.6 | 56.1 |

All but one of these preprint-journal article pairs were published prior to the current pandemic. However, comparing the results in tables 1 and 2, it becomes clear that turnaround times for articles that previously appeared as preprints are much shorter than the average turnaround times in these journals. In fact, for these pre-pandemic articles, turnaround times are even shorter than their post-pandemic counterparts.

## Conclusion

Our analysis indicates that the scholarly publishing enterprise has managed to drastically speed up the dissemination of Covid-19 related research material since the outbreak of the pandemic. In particular, academic journals managed to decrease the duration of their publication process by 49%, or 57 days on average, which is a statistically relevant difference. Some journals even show a decrease in publication time of over 80% compared to the pre-crisis era. This acceleration concerns both the stage of review (between submission and acceptance) and the editing stage (between acceptance and publication). The journals in our sample shortened both stages by 47% (45 days) and 61% (14 days) respectively. Hence the majority of the decrease in total publication time is due to speeding up the review process. We also conclude that the acceleration of the publication process is specific to Covid-19 related articles. Articles not related to Covid-19 published during the pandemic show very similar turnaround times as articles published before the pandemic.

In addition to a fast spread of information through journal articles, the number of papers submitted to preprint servers has drastically increased. However, such preprints do not seem to be taken up as journal publications any quicker than they were before the Covid-19 pandemic. Nevertheless, articles first appearing as preprints do seem to go through shorter publication processes than articles not appearing as preprints.

## Discussion

To tackle the current health crisis, many have urged to disseminate relevant academic knowledge as fast as possible. Acknowledging that typical publication delays in medical journals are unacceptable in the current era, journals are expected to decrease the turnaround times of their publication process. The results of our study indicate that journals have indeed managed to do so.

Our results on the average turnaround times of journal articles prior to the current pandemic corresponds well with earlier findings of studies on publication delays in medical journals (Björk & Solomon, 2013). However, since the outbreak of the Covid-19 pandemic, medical journals have managed to drastically accelerate their publication process to make it nearly twice as fast for Covid-19 related articles. On the contrary, articles not related to Covid-19 that were published since the beginning of the pandemic, do not show any acceleration. Their turnaround times are similar to articles published before the pandemic.

While it seems reasonable that journals might encounter difficulties to attract reviewers with relevant expertise – since those are probably active scientists working on novel research themselves – the contrary seems to be the case. Concluding from our results, it seems that journals are finding enough reviewers willing to review Covid-19 related papers on a very short notice. However, this conclusion should be treated with caution, as no data is available on *who* reviewed the papers. Maybe the same few experts reviewed a lot more than usual; maybe ‘relevant expertise’ was taken as a relative criterion, with journals using reviewers that usually would not have counted as experts. The fact that non-Covid-19 related papers are published at similar speeds during and before the pandemic, seems to indicate that journals are also not facing more issues with attracting reviewers for those papers.

As preprint articles are not being included in medical journals more quickly, it seems that either authors are not submitting preprint articles to journals more quickly, or journals are prioritizing content that has not appeared as preprint. Qualitative follow-up research interviewing authors and editors on their submission and review practices regarding preprint articles could shed further light on this.

Even though the acceleration of journals’ publication process is laudable from the perspective of quick information dissemination, it also raises several questions and concerns.

First, one could wonder whether faster is always better. Even though the two do not necessarily exclude each other, it seems reasonable that there is a balance, or perhaps even a trade-off, between speed and quality in peer review. Especially concerning the stage of review, legitimate concerns can be raised on whether speeding up the review process might harm the process’ ability to filter incorrect or invalid findings. Such research slipping through peer review, might require corrections or retractions in the future. Given the potentially rapid uptake of medical knowledge in policy and clinical contexts, such corrections might come in too late as potential harm might have already been done. Commentators have raised this concern regarding the usage of information in preprints, but it similarly applies to journal articles. In fact, false information spreading through journal articles is arguably more damaging, since it has the appearance of being ‘peer reviewed’ and hence properly verified. Scholars have repeatedly warned that a substantial share of articles (hastily) published during this crisis, will require future corrections (Marcus & Oransky, 2020). Formal expressions of concern – on papers used to make policy decisions – have already been issued (Voss, 2020). Future research should therefore analyse whether shorter review processes during the Covid-19 pandemic have led to an increase in corrections or retractions of published articles.

While drastic acceleration of the review stage might evoke quality issues, this arguably applies less to the editorial stage of the publication process. Journals’ achievement of shortening this stage of the publication process for Covid-19 related content is purely laudable. However, this might raise questions about why publication delays in this stage are usually higher and whether journals will aim and be able to maintain such standards in a post-crisis era. One potential explanation for the shortened editorial stage is that publishers or journal editors now prioritize Covid-19 related research articles, at the expense of other articles. However, our data on non-Covid-19 related articles published during the pandemic seems to contradict this. It seems like journals are managing to speed up editorial work for Covid-19 related content, while maintaining standards for other articles.

Several journals show a substantial lengthening of the editorial stage. This might be caused by an increase in the number of manuscripts submitted to the respective journals. For journals showing an increase in total turnaround time, this seems to be concentrated in the editorial process. Since editors themselves might be practicing scientists, the overload of newly submitted manuscripts might be a cause of this additional delay.

This study potentially suffers from various limitations in its analysis. First, it could only analyse those journal articles that have been published. This particularly implies that it was unable to assess the review process’ duration for rejected articles. Neither could it analyse articles that are currently still under review.

Second, the analysis does not include article type as a feature of analysis. Some article types, including letters to the editor, perspectives or commentaries, might undergo a different kind of peer review – they might for instance only be reviewed by the editor, rather than by external reviewers. A potential difference in distribution of pre- and post-crisis articles over the various article types might explain some of the variation in the publication process’ duration.

Third, our analyses focus on journals publishing relatively many Covid-19 related articles. Due to a lack of sufficient articles in other, potentially smaller journals, we were not able to analyse those journals. As larger journals may more easily attract reviewers and have more resources, and hence more capacity to shift resources, to execute the editorial stage of publication, the resulting decrease in publication time might be less strong in smaller journals. At a later stage, when more Covid-19 related papers appear in other journals, future research could verify this potential difference.

Last, it should be noted that some of the journal articles assigned to our control group concern papers related to previous health crises or pandemics such as the MESH, EBOLA or ZIKA crises. Despite similar incentives to publish those articles quickly, content related to Covid-19 makes its way through the publication process much quicker. Future research could include a more elaborate comparison not only between pre- and post-Covid-19 eras, but also between publishing in the Covid-19 and other health related crises.

In these times of crisis, the rapid dissemination of relevant academic knowledge is of paramount importance. Several stakeholders have already warned for a ‘fake-news pandemic’ spreading disinformation and conspiracy theories through social media channels in the absence of established scientific knowledge (Khatri et al., 2020; UNESCO, 2020). To assist policymakers and clinical experts, as well as to counter the spread of such disinformation, researchers and academic journals have a responsibility to share available knowledge quickly. The fact that medical journals have managed to considerably speed up their publication process for Covid-19 related content during the current pandemic is therefore laudable. However, some concerns remain about whether faster dissemination might go at the expense of research quality. Quickly spreading false information might do more harm than slowly spreading reliable knowledge.

## Acknowledgements

The author would like to thank the Centre for Science and Technology Studies at Leiden University for establishing the database used for this study and Digital Science for providing access to the data. The author also thanks Ludo Waltman, Nees Jan van Eck, Marc Luwel, Thed van Leeuwen and the members of the Institute for Science in Society’s Research Quality Team for valuable feedback on earlier drafts of this manuscript.

## References

Björk, B.-C., & Solomon, D. (2013). The publishing delay in scholarly peer-reviewed journals. Journal of Informetrics, 7(4), 914–923. doi:https://doi.org/10.1016/j.joi.2013.09.001

Brock, J. (2020). Rapid Registered Reports initiative aims to stop coronavirus researchers following false leads Nature Index.

Colavizza, G., Costas, R., Traag, V. A., van Eck, N. J., Van Leeuwen, T., & Waltman, L. (2020). A scientometric overview of CORD-19. bioRxiv.

Coronavirus pandemic: Nature’s pledge to you. (2020). Nature, 579, 471–472. doi:10.1038/d41586-020-00882-z

CWTS. (2020). CWTS_Covid Database. Retrieved from: https://github.com/CWTSLeiden/cwts_covid

Eisen, M. B., Akhmanova, A., Behrens, T. E., & Weigel, D. (2020). Publishing in the time of COVID-19. Elife, 9, e57162. doi:10.7554/eLife.57162

Gautret, P., Lagier, J.-C., Parola, P., Hoang, V. T., Meddeb, L., Mailhe, M., … Raoult, D. (2020). Hydroxychloroquine and azithromycin as a treatment of COVID-19: results of an open-label non-randomized clinical trial. International Journal of Antimicrobial Agents, 105949. doihttps://doi.org/10.1016/j.ijantimicag.2020.105949

Gunnarsdottir, K. (2005). Scientific journal publications: On the role of electronic preprint exchange in the distribution of scientific literature. Social Studies of Science, 35(4), 549–579.

Heimstädt, M. (2020). Between fast science and fake news: Preprint servers are political. Retrieved from https://blogs.lse.ac.uk/impactofsocialsciences/2020/04/03/between-fast-science-and-fake-news-preprint-servers-are-political/

Horbach, S. P. J. M., & Halffman, W. (2018). The changing forms and expectations of peer review. Research Integrity and Peer Review, 3(1), 8. doi:10.1186/s41073-018-0051-5

Horbach, S. P. J. M., & Halffman, W. (2019). The ability of different peer review procedures to flag problematic publications. Scientometrics, 118(1), 339–373. doi:10.1007/s11192-018-2969-2

Khatri, P., Singh, S. R., Belani, N. K., Yeong, Y. L., Lohan, R., Lim, Y. W., & Teo, W. Z. Y. (2020). YouTube as source of information on 2019 novel coronavirus outbreak: a cross sectional study of English and Mandarin content. Travel Medicine and Infectious Disease, 101636. doi:https://doi.org/10.1016/j.tmaid.2020.101636

Lin, Z., Hou, S., & Wu, J. (2016). The correlation between editorial delay and the ratio of highly cited papers in Nature, Science and Physical Review Letters. Scientometrics, 107(3), 1457–1464. doi:10.1007/s11192-016-1936-z

Marcus, A., & Oransky, I. (2020). The Science of This Pandemic Is Moving at Dangerous Speeds. Wired.

Nguyen, V. M., Haddaway, N. R., Gutowsky, L. F. G., Wilson, A. D. M., Gallagher, A. J., Donaldson, M. R., … Cooke, S. J. (2015). How Long Is Too Long in Contemporary Peer Review? Perspectives from Authors Publishing in Conservation Biology Journals. Plos One, 10(8), 20. doi:10.1371/journal.pone.0132557

Peters, D. P., & Ceci, S. J. (1982). Peer-review practices of psychological journals: The fate of published articles, submitted again. Behavioral and Brain Sciences, 5(2), 187–195. doi:10.1017/S0140525X00011183

Smith, R. (2006). Peer review: a flawed process at the heart of science and journals. Journal of the Royal Society of Medicine, 99(4), 178–182.

Talley, N. J. (2020). SARS-CoV-2, the medical profession, ventilator beds, and mortality predictions: personal reflections of an Australian clinician. n/a(n/a). doi:10.5694/mja2.50579

Teplitskiy, M., Acuna, D., Elamrani-Raoult, A., Körding, K., & Evans, J. (2018). The sociology of scientific validity: How professional networks shape judgement in peer review. Research Policy. doi:https://doi.org/10.1016/j.respol.2018.06.014

Thorp, H. H. (2020). Underpromise, overdeliver. 367(6485), 1405–1405. doi:10.1126/science.abb8492 %J Science

Tosi, H. (2009). It’s About Time!!!!:What to Do About Long Delays in the Review Process. 18(2), 175–178. doi:10.1177/1056492608330468

UNESCO. (2020). During this coronavirus pandemic, ‘fake news’ is putting lives at risk: UNESCO. Retrieved from https://news.un.org/en/story/2020/04/1061592

Voss, A. (2020). Statement on IJAA paper. Retrieved from https://www.isac.world/news-and-publications/official-isac-statement

