## Supplementary file A for "Pandemic Publishing: Medical journals drastically speed up their publication process for Covid-19"

**List of journals publishing most Covid-19 related articles**

| Journal title | number of articles | Articles in 2020 | Info on dates |
|---|---|---|---|
| Journal of Virology | 2485 | 4 | available |
| PLoS ONE | 1567 | 14 | available |
| Virology | 861 | 8 | available |
| Emerging Infectious Diseases | 810 | 72 | not available |
| The Lancet | 620 | 116 | not available |
| Viruses | 568 | 38 | available |
| Archives of Virology | 518 | 9 | available |
| Scientific Reports | 492 | 14 | available |
| Vaccine | 484 | 6 | available |
| Journal of Clinical Microbiology | 476 | 3 | available |
| Virus Research | 475 | 10 | available |
| Veterinary Microbiology | 444 | 8 | available |

Journals with data on dates available were selected.

**List of journals publishing most Covid-19 related articles since the outbreak of the pandemic**

| Journal title | number of articles | Articles in 2020 | Info on dates |
|---|---|---|---|
| The BMJ | 350 | 174 | not available |
| Science | 162 | 142 | not available |
| The Lancet | 620 | 116 | not available |
| Nature | 256 | 111 | not available |
| Journal of Medical Virology | 97 | 84 | available |
| The Lancet Infectious Diseases | 367 | 76 | not available |
| The New Scientist | 100 | 73 | not available |
| Emerging Infectious Diseases | 810 | 72 | not available |
| JAMA | 68 | 62 | not available |
| New England Journal of Medicine | 72 | 57 | not available |
| Case Medical Research | 56 | 56 | not available |
| Journal of Infection | 146 | 45 | not available |
| Travel Medicine and Infectious Disease | 121 | 45 | available |
| Eurosurveillance | 71 | 41 | available |
| Viruses | 568 | 38 | available |
| Chinese Medical Journal | 49 | 38 | not available |
| International Journal of Infectious Diseases | 260 | 33 | available |
| 中华结核和呼吸杂志 | 32 | 32 | not available |
| Journal of Hospital Infection | 172 | 26 | available |

Journals with data on dates available were selected.
